# Non-essential tRNA and rRNA modifications impact the bacterial response to sub-MIC antibiotic stress

**DOI:** 10.1101/2022.02.06.479318

**Authors:** Anamaria Babosan, Louna Fruchard, Evelyne Krin, André Carvalho, Didier Mazel, Zeynep Baharoglu

## Abstract

Antimicrobial resistance (AMR) develops as a major problem in infectious diseases treatment. While antibiotic resistance mechanisms are usually studied using lethal antibiotic doses, lower doses allowing bacterial growth are now considered as factors influencing the development and selection of resistance. Based on high throughput transposon insertion sequencing (TN-seq) in *V. cholerae*, we have undertaken the phenotypic characterization of 23 transfer RNA (tRNA) and ribosomal RNA (rRNA) modifications deletion mutants, for which growth is globally not affected in the absence of stress. We uncover a specific involvement of different RNA modification genes in the response to aminoglycosides (tobramycin (TOB), gentamicin (GEN)), fluoroquinolones (ciprofloxacin (CIP)), β-lactams (carbenicillin (CRB)), chloramphenicol (CM) and trimethoprim (TRM). Our results identify t/rRNA modification genes, not previously associated to any antibiotic resistance phenotype, as important factors affecting the bacterial response to sub-MIC antibiotics from different families. This suggests differential translation and codon decoding as critical factors involved in the bacterial response to stress.

## Introduction

Antibiotic overuse and misuse contribute to AMR, via selective pressure exerted by treatment during infection, but also in the environment where gradients of antibiotics are found in soil and water, the natural reservoir of many bacteria among which *Vibrios*. AMR is increasingly associated with life in the aquatic environment, particularly in aquaculture farms, where several bacterial species coexist. A World Health Organisation report on AMR in enteric pathogens states that “consideration must be given to the relationship of *Vibrios* with the environment” to understand AMR development^1^. Most studies address the bacterial response to lethal antibiotic concentrations and the effect of gene mutations on antibiotic resistance. Meanwhile, in their environments, bacteria encounter sub-minimal inhibitory concentrations (sub-MICs) of antibiotics^2^, which are stressors, and can lead to transient phenotypic tolerance to high doses of antibiotics^3^. Thus, characterization of the bacterial responses to such stress and its impact on resistance/tolerance, need to be comprehensively clarified.

We have previously demonstrated that several pathways identified for the response to antibiotic stress in *V. cholerae* are paradigmatic for other bacterial pathogens^4,5^. Using sub-MIC antibiotics, we aimed at characterizing which bacterial responses were triggered and allowed the cells to grow and survive, and we asked whether the identified processes also impact bacterial phenotypes at lethal concentrations of the same antibiotics. Our results point to a central role of transfer RNA (tRNA) and ribosomal RNA (rRNA) modifications in the response to sub-MIC antibiotic stress, suggesting that RNA modification profiles and translation may be modified in bacteria by stress.

Evolution of resistance requires genetic diversity in populations, yet non-genetic phenotypic diversity can also contribute. One process generating phenotypic diversity is translation, with an error rate up to 10^−3^ substitutions per position^6^. Translation errors cause protein misfolding^7,8^, aggregation and proteotoxic stress^9^. Translation errors can also provide transient increase in fitness^10^, offering cells the necessary time to acquire beneficial genetic mutations^11^ and to eliminate deleterious ones^12^, as it was observed upon oxidative stress^13^ and proteotoxic stress^14^. One can thus presume a tradeoff between overabundant errors causing toxicity under favorable growth conditions and insufficient errors hampering survival in conditions of stress.

Codon decoding efficiency can impact translation speed or translation accuracy at specific mRNAs/codons and proteome diversity^6^. Differences in decoding and reading frame maintenance have already been linked with the presence or absence of certain RNA modifications^15,16^. In particular, methylation at specific positions in rRNA stabilizes the binding of initiator tRNA to the ribosome at the start codon^17^, and several rRNA methylation factors have been linked to AG resistance^18,19^. Regarding tRNA modifications, more than 80 have been described in bacteria^20^. They can be involved in tRNA stability^21^, abundance^6^, decay^22,23^ and affinity for the ribosome^24^. While some tRNA modification genes are essential, (e.g. *trmD, tadA*), in many cases their deletion does not confer any visible phenotype to the unstressed cells^20^ (**Table S1** and references therein). Few studies address the exact physiological roles of non-essential rRNA^25,26^ and tRNA modifications in bacterial stress response phenotypes (^27-32^, reviewed in^20^).

In the present study, we link antibiotic stress with RNA modification genes different from previously known ones. We show that their inactivation confers, not resistance, but increased or decreased fitness in presence of antibiotic stress.

## Results

### TN-seq identifies rRNA and tRNA modification genes involved in the response to sub-MIC TOB and CIP in *V. cholerae*

Using TNseq in *V. cholerae*, we searched for genes that are important for growth in the presence of sub-MICs of antibiotics targeting the ribosome (TOB belonging to aminoglycosides (AGs)), or DNA (CIP, belonging to fluoroquinolones (FQs)). We constructed large transposon inactivation libraries in *V. cholerae* as previously performed^33^ and we subjected them to growth without or with antibiotics at 50% of the minimal inhibitory concentration (MIC), during 16 generations. After sequencing and bioinformatics analysis of the regions flanking the transposon, we identified genes where reads associated to detected transposon insertions increase or decrease. Loss of detected insertions in a specific gene generally means that the inactivation of this gene is detrimental in the tested condition, while enrichment means that the inactivation is beneficial. In some cases, transposon insertion in one gene may also lead to differential expression of downstream genes. In this study, we searched for genes that are important only during sub-MIC treatment. We thus compared insertion counts after 16 generations in sub-MIC antibiotics (TOB or CIP) to those after 16 generations without antibiotics (**Figure 1** and **Tables 1 and S1**). Genes having a significant impact on fitness (insertions enriched or lost) in the non-treated condition are thus not included in our analysis. For both antibiotics, we found common or antibiotic specific RNA modification genes whose number of reads was impacted, suggesting that their inactivation was either beneficial or detrimental for growth in the presence of the sub-MIC antibiotic.

**Figure 1.**
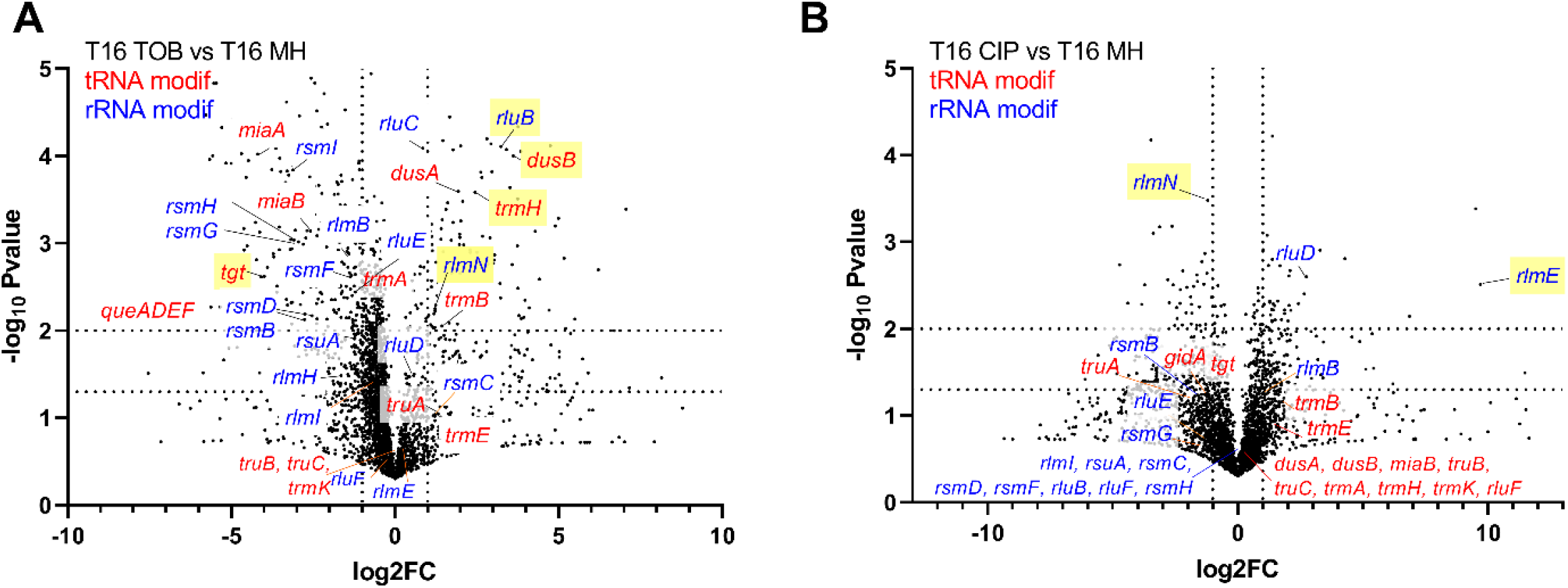
TN-seq identifies rRNA and tRNA modification genes affecting fitness of *V. cholerae* in the presence of sub-MIC TOB and CIP. AB: tRNA modification genes are indicated in red. rRNA modification genes are indicated in blue. *rlmN* modifies both tRNAs and rRNA. Volcano plots represent genes for which transposon inactivation is beneficial or detrimental after 16 generations of growth, compared to growth without antibiotics. A: TOB 50% of the MIC, B: CIP 50% of the MIC. X axis represents log2 fold change of the number of transposon reads associated with gene inactivations, detected after 16 generations in the indicated antibiotic versus non-treated condition. The Y-axis represents the negative log_10_ *p value*. Gene inactivations which show the strongest antibiotic specific effects are highlighted in yellow.

**Table 1.**
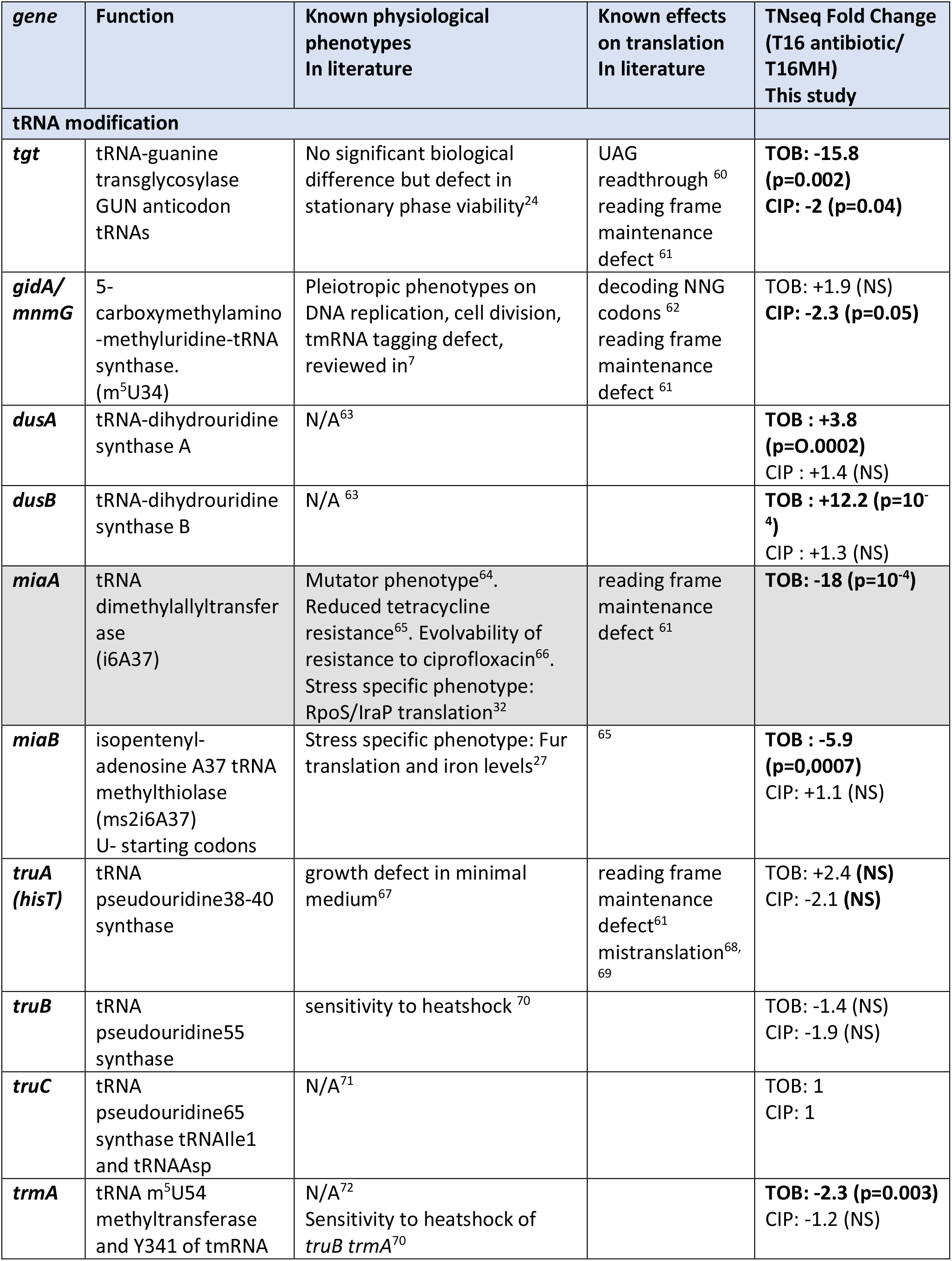
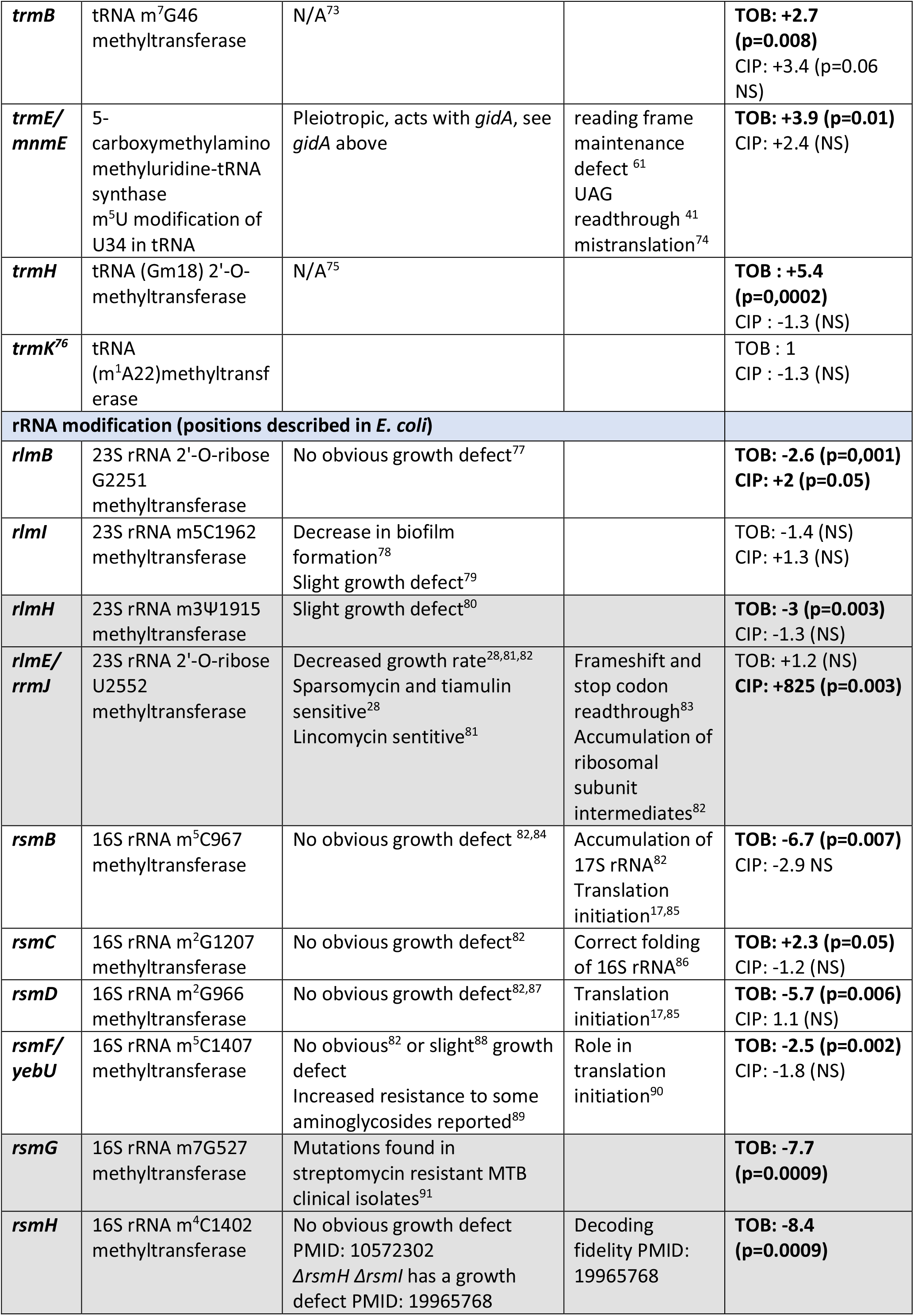
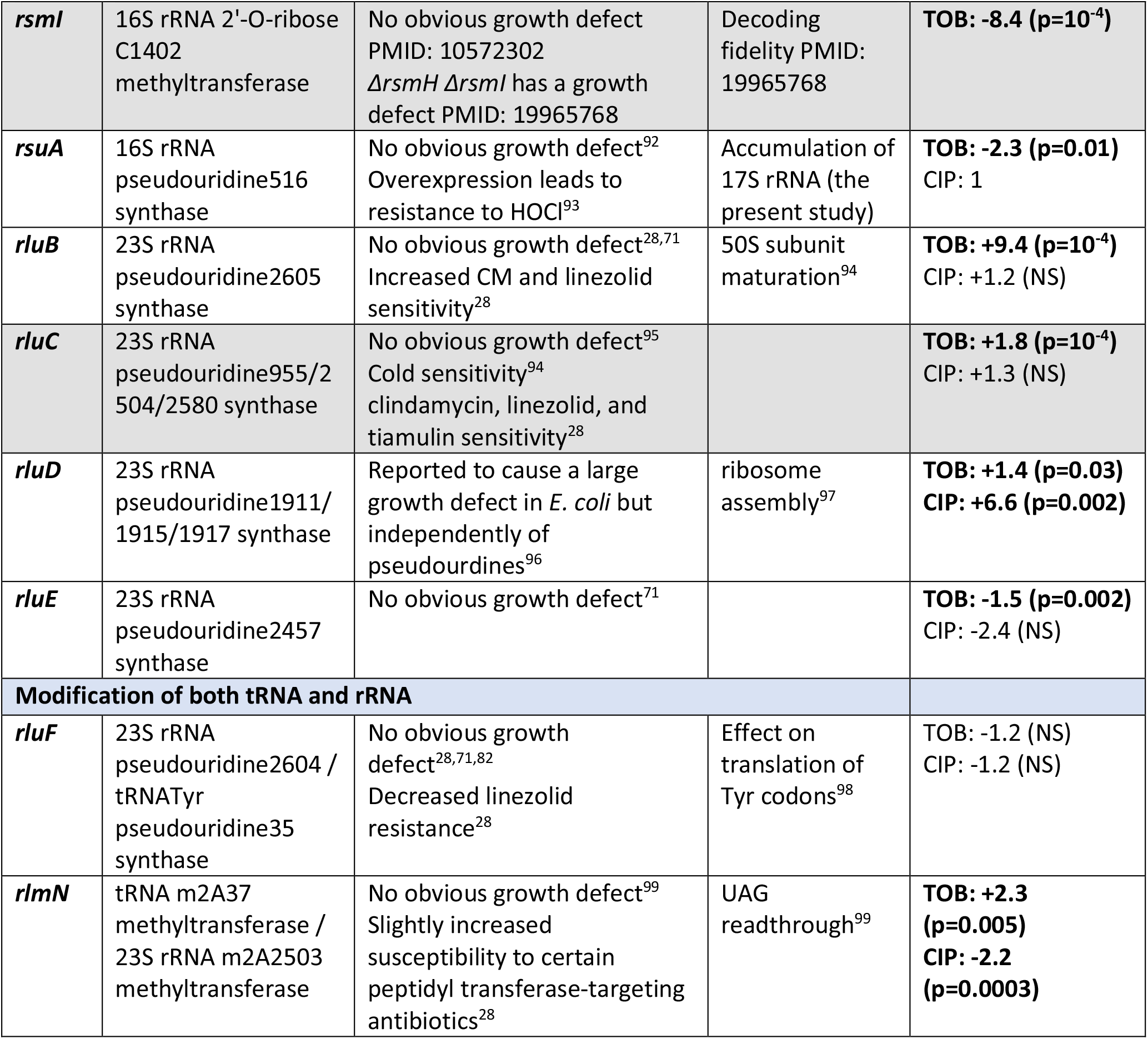
Phenotypes associated to RNA modification genes identified by TNseq. In grey: not selected for further study. NS : non-significant p value.

The most important TNseq hits for TOB include: (i) tRNA modification genes for which inactivation is detrimental: incorporation of queuosine (Q) by *tgt* (together with the Q synthesis genes *queADEF)*, and i6A37/ms2i6A37 by *miaA/miaB*; or beneficial: dihydrouridine (D) incorporation by *dusB, dusA*; and methylation by *trmH, rlmN*; and m^5^U34 incorporation (*gidA*, also called *mnmG*); (ii) rRNA modifications for which inactivation is detrimental: methylation by *rsmI, rsmF, rsmG, rsmH, rsmB, rsmD*, pseudouridine (ψ) incorporation by *rsuA* ; or beneficial: ψ by *rluB*. Note that *rsmG* and *rsmF* mutants have already been associated with increased AG resistance (**Table 1** and references therein), but our results suggest decreased fitness in AGs for these mutants.

For CIP: (i) tRNA modification genes for which inactivation is detrimental were responsible for ψ incorporation (*truA)* and methylation (*rlmN)*; (ii) rRNA methylation genes were also identified in CIP, some at different positions than those in TOB (detrimental inactivation of *rsmB* and beneficial *rlmE*). Note that RlmN can modify both tRNA and rRNA.

Overall, several non-trivial observations stem from our results: first, the effect of inactivation of these genes on fitness can either be negative (e.g. *tgt* in TOB), or positive (e.g. *dusB* in TOB). Second, their impact seems to be an antibiotic specific one. For instance, the inactivation of *dusB*/*tgt*/*rluB* strongly impacts the fitness in TOB, but not in CIP; and inactivation of *rlmN/gidA/rlmB* even affect fitness in opposing ways in TOB and CIP.

These observations suggest that the loss of a given modification may affect the bacterial response in a specific way rather than through a general effect of all modifications on translation. While AGs, which target the ribosome, could be expected to impact translation related genes, it was surprising that the response to CIP which targets DNA also involves several RNA modification genes, suggesting that the involvement of RNA modifications may be fundamental upon stress due to antibiotics from different families.

### RNA modification gene deletions impact fitness during growth in sub-MIC antibiotics

We next constructed *V. cholerae* deletion mutants for 23 of the identified RNA modification genes, selected in TNseq data for having no (or slight) effect on fitness during growth in the absence of antibiotics. Many have no known physiological defect, and were not previously associated to antibiotic related phenotypes (**Table 1**). The following genes were excluded from further study either for known effects on growth: *miaA, rsmA, rlmE;* or for known AG related phenotypes: *rsmG, rsmA* & *rsmH*^26^. We chose *trmK* as a neutral control for TOB and CIP, as it was not identified in our TNseq screens.

Since growth curves of monocultures of the mutants were similar to that of the WT in the absence of treatment (not shown), we decided to perform competition experiments between mutants and the WT strain, to assess effects on fitness in sub-MICs of 6 different antibiotics: the AGs TOB and gentamicin (GEN), the fluoroquinolone CIP, as used in our TNseq screen, and additionally the β-lactam carbenicillin (CRB) targeting the cell envelope, chloramphenicol (CM) targeting translation elongation and trimethoprim (TRM) which inhibits thymidine synthesis interfering with DNA synthesis. **Figure 2** shows the competitive index of mutants compared to WT.

**Figure 2.**
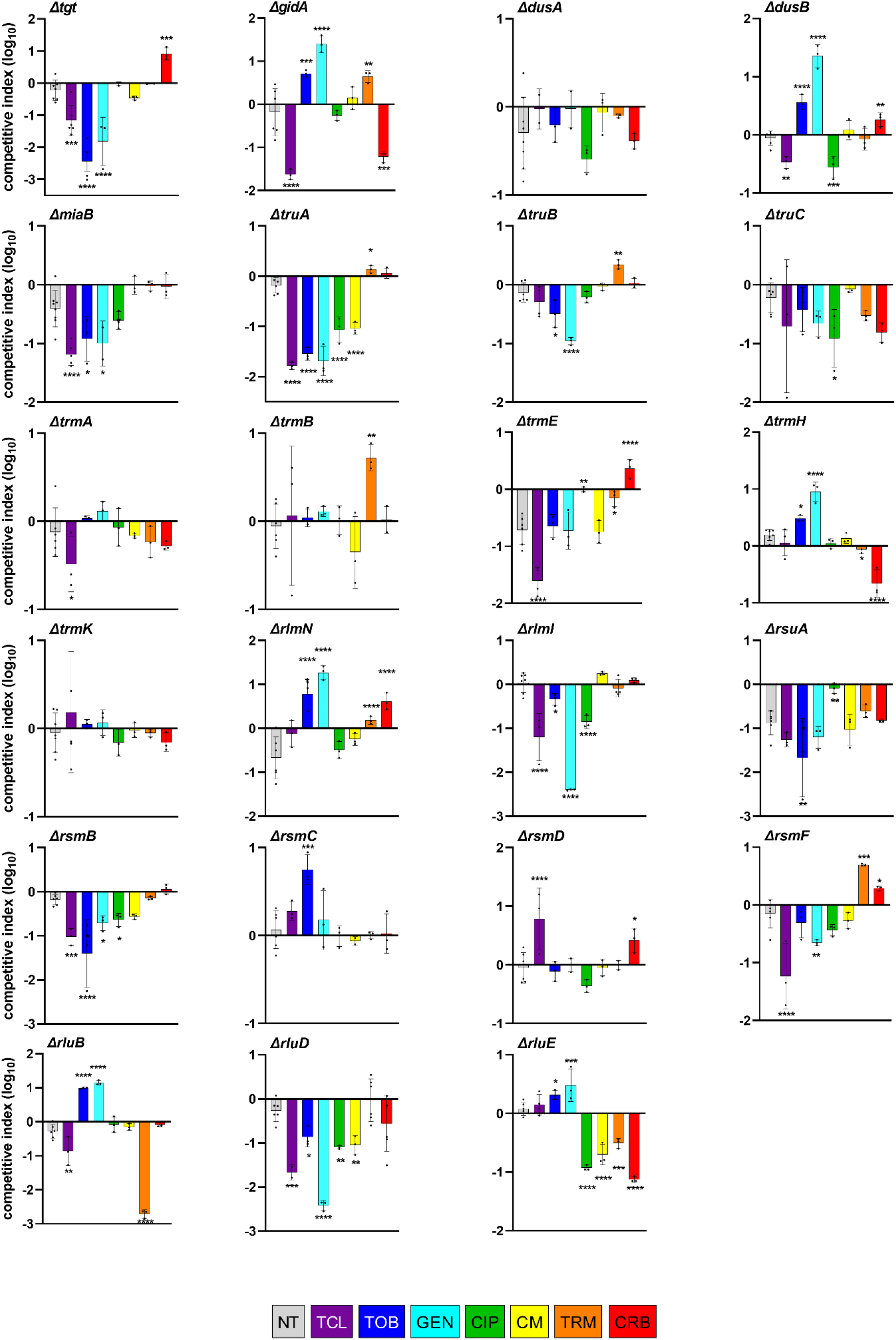
Impact of RNA modification gene deletions on fitness during growth in sub-MIC antibiotics. In vitro competition experiments of *V. cholerae* WT and mutant strains in the absence or presence of different antibiotics at sub-MICs (50% of the MIC, TCL: triclosan 0.01 mM, TOB: tobramycin 0.6 μg/ml; GEN: gentamicin 0.5 μg/ml; CIP: ciprofloxacin 0.002 µg/ml, CM: chloramphenicol 0.4 µg/ml, TRM: trimethoprim 0.4 µg/ml, CRB: carbenicillin 2.5 µg/ml). The Y-axis represents log_10_ of competitive index calculated as described in the methods. A competitive index of 1 indicates equal growth of both strains. NT: no antibiotic treatment. For multiple comparisons, we used one-way ANOVA on GraphPad Prism. **** means p<0.0001, *** means p<0.001, ** means p<0.01, * means p<0.05. Number of replicates for each experiment: 3<n<8.

As expected, deletions of the majority of tested genes (with the exception of *trmE, rsuA* and *rlmN*) have no or little effect on competitive index during growth in the absence of antibiotics (**Figure 2**), emphasizing their specific role during stress, here sub-MIC antibiotics.

For the AGs TOB and GEN, among tested genes, deletion of *tgt, miaB, truA, truB, rlmI, rsmB, rsmF, rluD* decreased fitness, while deletion of *gidA, dusB, trmH, rlmN, rsmC, rluB, rluE* conferred a growth advantage. These results were consistent with TNseq data, with the exception of *truA, truB, rluE, and rluD* for which the TNseq data were not statistically significant. For CIP, deletions of *dusB, miaB, truA, truC, rlmI, rsmB, rluD, rluE* were disadvantageous, whereas *ΔtrmE* and *ΔrsuA* strains appear to lose their fitness disadvantage compared to WT. Once again, results were consistent with statistically significant TNseq results, except for the *rluD* gene. For CM, *truA, rluD* and *rluE* deletions were detrimental. For TRM, *rluB* and *rluE* deletions were detrimental, while deletions of *gidA, truB, trmB, rlmN, rsmF* conferred a low (up to 10x) but statistically significant growth advantage. For CRB, detrimental deletions were *gidA, trmH, rluE*, and advantageous deletions were *tgt, dusB, trmE, rlmN, rsmD, rsmF*.

In order to test whether these modification genes could be important for the response to another type of stress, we also performed competitions in the presence of the biocide triclosan (TCL), at 50% of the MIC. TCL inhibits fatty acid synthesis and can be found in antiseptic consumer products. It has been a subject of concern for its impact on the aquatic environment^34^ and antibiotic resistance development^35^. Again, while deletion of many RNA modification genes decreased fitness in TCL (*tgt, gidA, dusB, miaB, truA, trmA, trmE, rlmI, rsmB, rsmF, rluB, rluD*), some were neutral (*dusA, trmB, trmH, rlmN, rluE, trmK*), and one was beneficial (*rsmD*).

These results globally confirm that the effect of a given modification gene is not a general one on viability but an antibiotic specific one. For instance, regarding tRNA modifications, upon AG treatment (TOB, GEN), deletion of *tgt* confers a clear 10 to 1000x disadvantage, while it has no major effect in CIP, TRM, CM, MMC, and appears to be 10x advantageous in CRB. Deletion of *truA* confers a up to 100x fitness defect in AGs, CIP and CM but is neutral in TRM and CRB. Deletion of *truB* also appears to affect specifically growth in AGs. Deletions of *dusB/rlmN*, and *gidA*/*trmH* are highly (10 to 100x) beneficial in AGs but respectively deleterious or neutral in CIP. *rlmN* deletion also confers a slight advantage in TRMand CRB. Deletion of *trmA* shows no major effect in any antibiotics, while *trmB* deletion is only beneficial in TRM. Regarding rRNA modifications, *rluB* shows a striking phenotype with 10x beneficial deletion in AGs, highly (1000x) deleterious in TRM, and neutral in presence of the other antibiotics. Of note, *gidA* (*mnmG)/ trmE (mnmE)* are known to have pleiotropic phenotypes due to effects on translation^36^, chromosome replication and cell division^20,37-40^, in addition to effects on tRNA modification^41^. Regarding TCL, many RNA modification gene deletions confer a fitness defect. However, the fact that deletion of *rsmD* is beneficial indicates that bacteria can also have an active response mechanism to the presence of toxic chemicals such as antiseptics.

### RNA modification gene deletions impact tolerance to high doses of antibiotics without changing the resistance

Next, we addressed whether these genes could be involved in the response to lethal antibiotic concentrations. We first determined the minimal inhibitory concentrations (MIC) of TOB, CIP, TRM and CRB for each deletion mutant (**Table S2**). Slight decreases in the MIC of TOB, in the order of 10%, was observed for *ΔrlmI* and *ΔrsmD*. Slight increases in MIC were observed for *ΔgidA* and *ΔrluB* in TOB (x1,6), for *ΔgidA, ΔrluD, ΔtrmE* in CIP (x1,2), (x1,1) for *ΔrlmN* in ampicillin (as a substitute for CRB) and for *ΔtrmE* and *ΔtruC* in TRM (x1,6). Besides these small changes, we found no major differences in MICs, consistent with the fact that these genes were not previously associated with antibiotic resistance phenotypes.

We then tested the survival to lethal concentrations of antibiotic (**Figure 3**): WT and mutant bacteria were grown to early exponential phase and then treated for 20 hours with 10xMIC of TOB, CIP, TRM and CRB as previously performed^42^.

**Figure 3:**
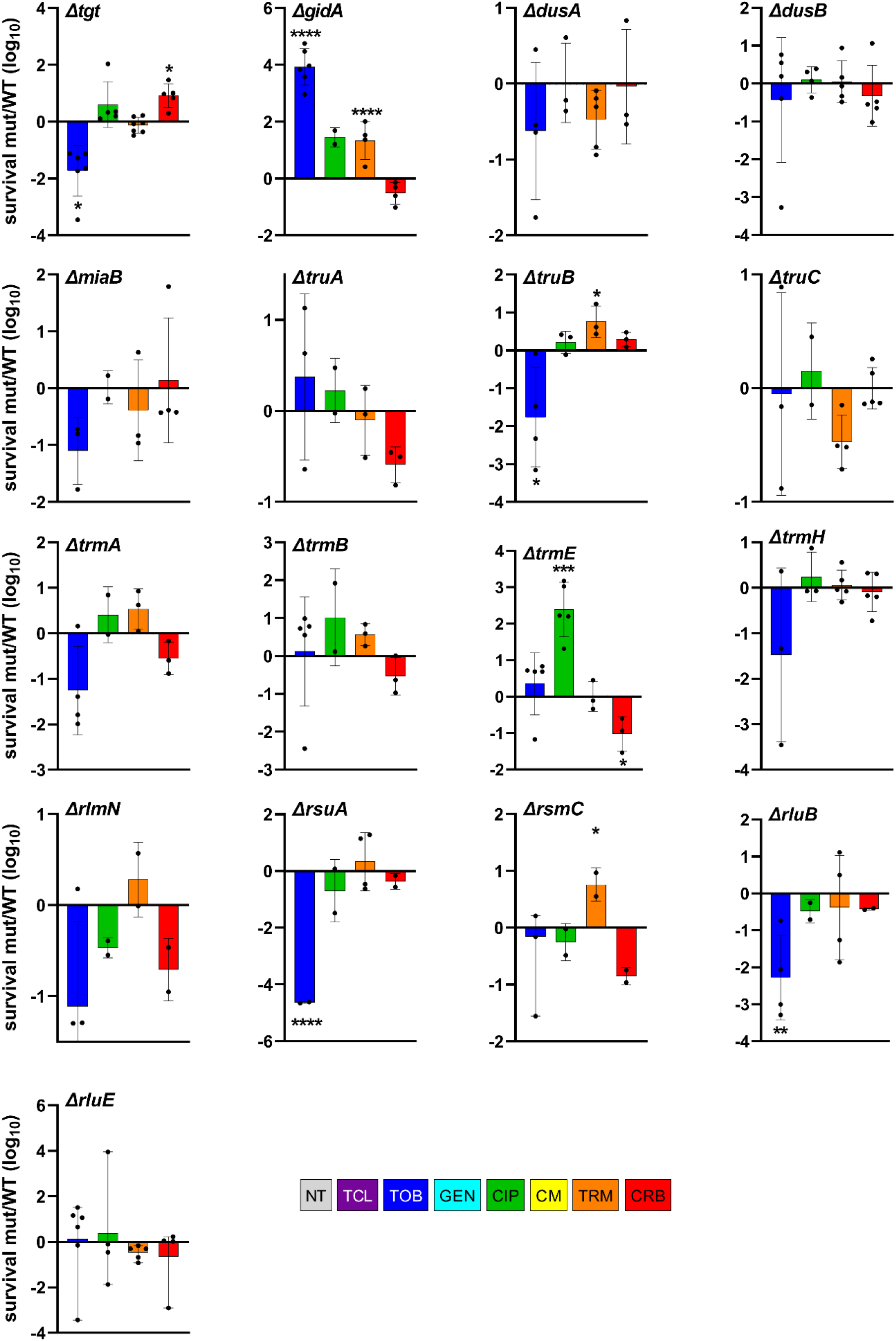
Survival to lethal antibiotic treatment. *V. cholerae* WT and deletion mutant cultures were grown without antibiotics up to early exponential phase. Total number of bacteria (T0) was determined by plating on MH plates before addition of the indicated antibiotic at >MIC, at time T0. After 20 hours incubation with the antibiotic, the number of surviving bacteria was determined and plating on MH plates (T20). Survival was calculated for each strain by dividing the number of surviving bacteria at T20 by the initial number of bacteria. The Y-axis represents the log10 survival ratio of a given mutant over the survival of the WT strain. Antibiotic concentrations: tobramycin 10 µg/ml, ciprofloxacin 0.04 µg/ml, trimethoprim 50 µg/ml, carbenicillin 50 µg/ml. Means and geometric means for logarithmic values were calculated using GraphPad Prism. For multiple comparisons, we used one-way ANOVA on GraphPad Prism. **** means p<0.0001, *** means p<0.001, ** means p<0.01, * means p<0.05. Number of replicates for each experiment: 3<n<8.

For 10 mutants out of 17 tested (among which 9 tRNA mutants), survival profiles were consistent with fitness profiles shown in **Figure 2**. These were mutants *tgt, gidA, truB, trmE* (except in CRB) and *rsuA*, for which increased fitness corresponded to increased tolerance and vice-versa; and *dusA, miaB, truC, trmA, trmB* for which the absence of statistically significant effect on tolerance was also consistent with the absence of differences in fitness. This suggest that a fitness (dis)advantage in sub-MIC antibiotics in the absence of tRNA (and rRNA) modifications may also impact tolerance to lethal doses of the same antibiotic, without changing the MIC.

For 3 mutants, *dusB, trmH, rluE*, no significant effect on tolerance was generally observed at 20h of lethal treatment, while deletion of these genes positively affected fitness in sub-MIC TOB. In order to address whether differences in tolerance could be detected at earlier times of treatment, we repeated the experiments and spotted cultures after 30 min, 1 and 2 hours of antibiotic treatment instead of 20 hours **(figure S1**). While *ΔdusB* tolerance was still similar to that of WT, *ΔtrmH* and *ΔrluE* strains displayed increased tolerance to TOB at 30min and 1h of treatment, consistent with their beneficial effect on fitness in sub-MIC TOB.

For the remaining 4 mutants, among which 3 rRNA modification mutants, we observed contradictory phenotypes between fitness and 20h tolerance, i.e. decreased TOB tolerance at 20h in beneficial deletion mutants *rlmN, rsmC, rluB;* and in CRB for *trmE*. However, at earlier time point as described above, and consistent with fitness profiles, TOB tolerance is clearly increased in *rlmN, rsmC, rluB* (**figure S1**), suggesting that mutants with fitness advantage in sub-MIC TOB also survive longer in the presence of lethal TOB concentrations. However, the final survival after 20h of treatment is not increased, consistent with unchanged MICs. This phenotype is a characteristic of antibiotic tolerant populations^43^. CRB tolerance of *ΔtrmE* remains lower than WT (not shown).

Overall, tolerance profiles of several mutants correlate with their fitness profiles in sub-MICs of antibiotics. For those, such as *ΔdusB*, with increased fitness but not tolerance, the mechanisms remain to be determined, and their phenotypes suggest that the effects of RNA modifications during growth in stressful (sub-MIC antibiotic) conditions do not necessarily affect survival to high antibiotic doses. rRNA modifications in particular could be expected to have structural effects on ribosomes, which could lead to pleiotropic effects, and could potentially explain this discrepancy.

One such effect is 17S rRNA accumulation, due to a defect of maturation to 16S rRNA (previously shown for *ΔrsmA*^44^ and *ΔrsmB*, **Table 1**). We visualized rRNA species purified form exponentially growing WT and RNA modification deletion mutants (**Figure S2**). We find accumulation of a pre-16S, consistent with 17S, rRNA species for *ΔrsuA*, for which fitness is most affected also in the absence of antibiotics. RsuA is a 16S rRNA pseudouridine synthase. Apart from *ΔrsuA*, we observed no differences in rRNA species between the other tested deletion mutants and the WT. This is consistent with the fact that these strains do not exhibit any major growth defect in the absence of antibiotics. Further study is needed to clarify the role of identified rRNA modifications in antibiotic specific survival.

We also evaluated whether deletion of these RNA modification genes could have any effect on DNA mutation rates by quantifying the appearance of spontaneous rifampicin resistant mutants (**Figure S3**), and found no major effect on mutation rates except for *ΔgidA*. This confirms that the fitness advantage/disadvantage conferred by RNA modification gene deletions are not due to an effect on mutation rates and/or accumulation of mutations.

### RNA modification gene deletions also impact *E. coli* growth in sub-MIC antibiotics

We next sought to determine whether RNA modification genes also play similar roles in bacterial species other than *V. cholerae*. We constructed deletion mutants in *E. coli* MG1655 of 9 genes selected for their positive (*gidA, dusB, rsmC, rluB*), neutral (*dusA, rsmD*) and negative (*tgt, trmE, rsuA*) impact on *V. cholerae* fitness in sub-MIC TOB (**Figure 4** and **Figure S4**). Note that inactivation of *dusA* and *rsmD* were observed to be respectively beneficial and deleterious in *V. cholerae* TN-seq data, but not in competitions. Growth curves in 50% MIC TOB display similar and dissimilar phenotypes in *E. coli* compared to those observed for *V. cholerae*. First, similar to *V. cholerae*, (i) deletions of *dusB, rsmC, rluB* and *dusA* and (ii) deletions of *tgt, trmE* and *rsmD*, respectively have a positive and a negative impact on growth in sub-MIC TOB in *E. coli*. For *Δtgt*, we also had some replicates with no observable effect in sub-MIC TOB (curve in light blue), suggesting heterogeneous response to TOB stress in this mutant in *E. coli*. On the other hand, unlike in *V. cholerae, ΔgidA* decreases while *ΔrsuA* improves growth of *E. coli* MG1655 in TOB. Note that synteny is conserved between the two organisms for these genes, hence the differences cannot be attributed to an effect of the deletions on surrounding genes. We also observed the same growth profiles (**Figure S5**) in an *E. coli* BW25113 (Keio) strain, for all mutants except for the BW25113 *Δtgt* strain which unexpectedly has a positive impact on growth in sub-MIC TOB in this genetic context. Note that growth curves show differences in growth but not necessarily in fitness as it is the case for competition experiments where WT and mutant cultures are mixed. Nonetheless, results show that the involvement of RNA modification genes in the response to sub-MIC antibiotic stress is not specific to *V. cholerae* and can be extended to other bacterial species, although their antibiotic related effects may sometimes be species and even strain-specific.

**Figure 4.**
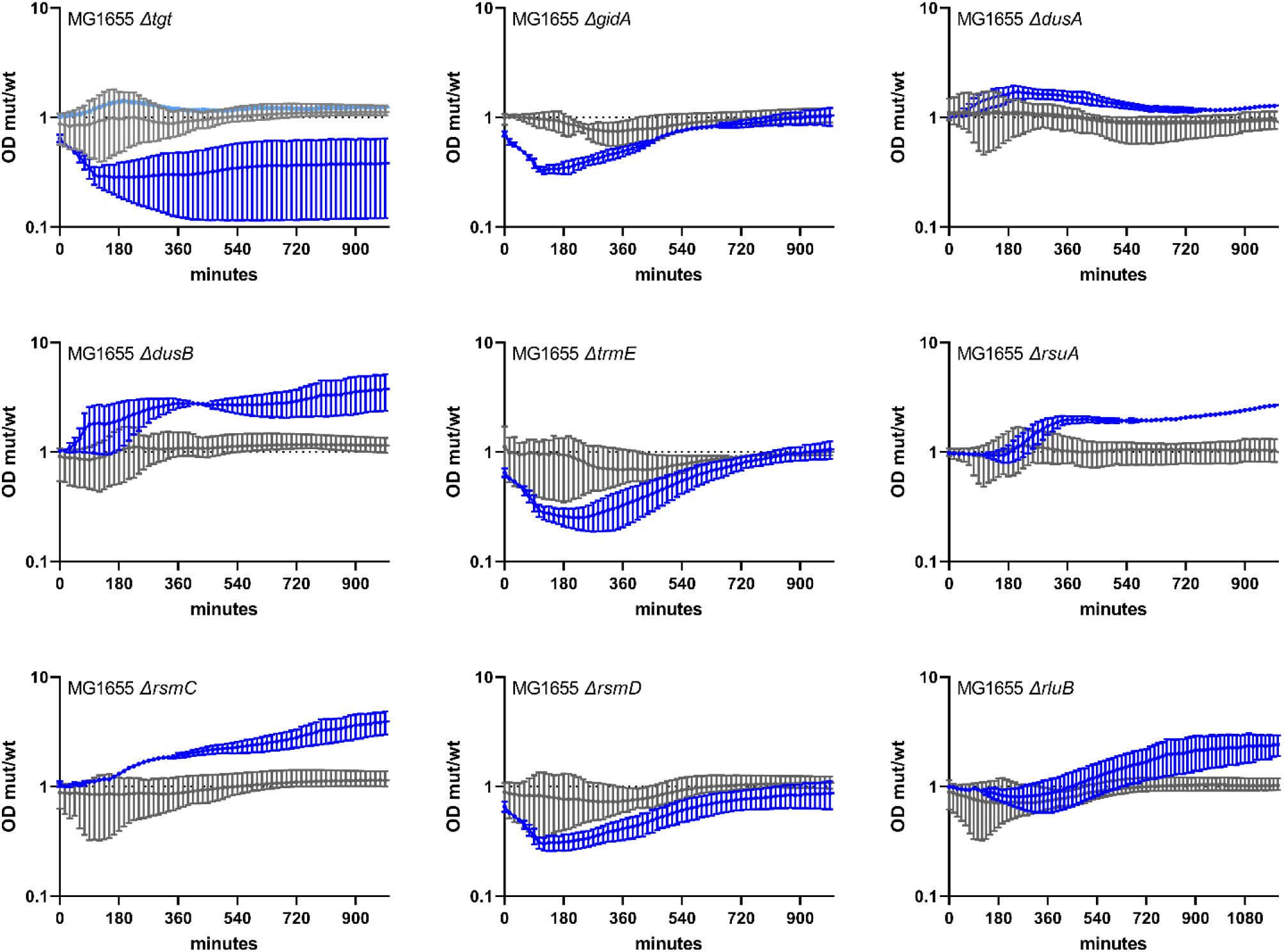
Growth of *E. coli* MG1655 WT and derivatives deleted for selected RNA modification genes in sub-MIC TOB. Overnight cultures were diluted 100x in fresh MH medium, on 96 well plates. Each well contained 200 µl. Plates were incubated with shaking in TECAN plate reader device at 37°C, OD 620 nm was measured every 15 minutes. Grey: no treatment. Blue: sub-MIC TOB, at 0.2 µg/ml (50% of the MIC for MG1655 in MH liquid culture). The Y-axis represents the OD of the mutant divided by the OD of the WT strain in the same growth conditions, and thus reflects slower (below 1) or faster (above 1) growth. Standard deviation is shown.

## Discussion

Using antibiotics at sub-MICs, we identify here the importance of rRNA and tRNA modification genes, not previously associated to any antibiotic resistance/tolerance phenotypes (**Table S1** and references therein). Among these are rRNA methylation factors RsmB/C/D/H/I and RlmI, rRNA pseudouridine synthases RsuA and RluD; and Tgt, DusB, TruA/B/C, TrmA/B/E/H and RlmN for tRNA modifications.

Most t/rRNA modifications influence translation rate, fidelity and precision of codon decoding^45^. Errors in decoding can lead to transient tolerance to stress^10^, increasing the cell’s chances to acquire genetic mutations allowing adaptation^12^. For instance, increased survival after 20h antibiotic treatment as described above for several mutants, may be due to tolerance or persistence, a transient state of phenotypic (non-heritable) resistance to lethal ATB concentrations. The idea that RNA modifications can act on such phenotypic adaptation is interesting, and worth pursuing.

Since the genetic code is degenerate, and one tRNA can decode several codons, decoding efficiency can logically be impacted by tRNA modification^46^. Thus, the link between tRNA modification-dependent differences in translation, proteome diversity, and the bacterial response to antibiotics and more generally to changing environments, is an attractive area to explore further. It is known that codon usage has an impact on translation^47-49^. Highly translated mRNAs, such as those of ribosomal proteins, display a codon usage profile different than the general codon usage in the genome^50^. It was proposed that codon usage of highly expressed genes is determined by the abundance of tRNAs, so as to prevent titration of tRNAs, hence allowing efficient translation of the rest of the proteome^51^. We can speculate that codon usage of these genes can also be a function of decoding efficiency displayed by modified vs. unmodified tRNAs. Various transcriptional regulators also show codon usage biases, and RNA modifications may impact their translation and thus lead to differential transcription of the regulon that they control^27,29,31,32^.

Stress regulated RNA modifications would facilitate homeostasis by reprogramming the translation of stress response genes^52^. Although RNA modifications were initially thought to be static, studies reveal the existence of dynamic modifications depending on growth phase and rate^53^, environmental changes (reviewed in^25,54^) or stress^55^, leading to differential translation of stress response transcripts and translational reprogramming^52^. In this process, RNA modifications and modification levels have an impact on the translation of regulators, which were thus defined as modification tunable transcripts, or MoTTs^56^.

Such processes were described in *E. coli* for the general stress sigma factor *rpoS* carrying leucine codons necessitating MiaA-modified tRNAs^32^; for the iron sensing *fur* regulator, carrying serine codons decoded by MiaB-modified tRNA, in response to low iron^27^; for the response to magnesium levels through TrmD modification dependent decoding of proline codons in *mgtA*^30^. In *Pseudomonas aeruginosa*, TrmB modification increases translation efficiency of phenylalanine and aspartate enriched catalase mRNAs during oxidative stress^29^, suggesting tRNA methylation mediated translational response to H_2_O_2_. During the mycobacterial response to hypoxic stress^31^, differential translation of specific stress response genes was linked, first *in silico*, then experimentally, to their codon usage bias. Our results highlight tRNA dependent translational reprogramming as a promising subject to be addressed in bacteria in regard to antibiotic stress.

This study reveals the existence of an epigenetic control of the response to sub-MIC antibiotics at the RNA level, adding upon our previous report of an epigenetic tolerance to aminoglycosides at DNA level^57^. Such a response may also involve gene sequences which co-evolve with the specific bacterial species so that translational regulation of the response to antibiotics becomes associated with other stress response genes bearing differentially decoded sequences, i.e. modification tunable transcripts. It can also not be excluded that certain of these RNA modification enzymes also exert their effect through mRNA modification^58,59^. Molecular study of codon decoding particularities of each RNA modification mutant, coupled to proteomics and *in silico* analysis of genes with differential codon usage, can allow for the identification of specific pathways post-transcriptionally regulated by a given RNA modification.

## Acknowledgements

We thank Manon Lang for her warm support with the setting up of survival and molecular biology experiments, Chloé Korlowski for assistance with *E. coli* deletion strain constructions and Sebastian Aguilar-Pierlé for assistance with TNseq library analysis. This research was funded by the Institut Pasteur, the Centre National de la Recherche Scientifique (CNRS-UMR 3525), ANR ModRNAntibio (ANR-21-CE35-0012), the Fondation pour la Recherche Médicale (FRM EQU202103012569) and Institut Pasteur grant PTR 245-19. AB was funded by Institut Pasteur Roux-Cantarini fellowship.

---

**Table 1. Phenotypes associated to RNA modification genes identified by TNseq**. In grey: not selected for further study.

**Table S1. TNseq analysis for the whole genome**.

**Table S2. Minimal inhibitory concentrations determined using *etests***.

**Table S3. Strains and plasmids**.

**Table S4. Primers**

**Figure S1:**
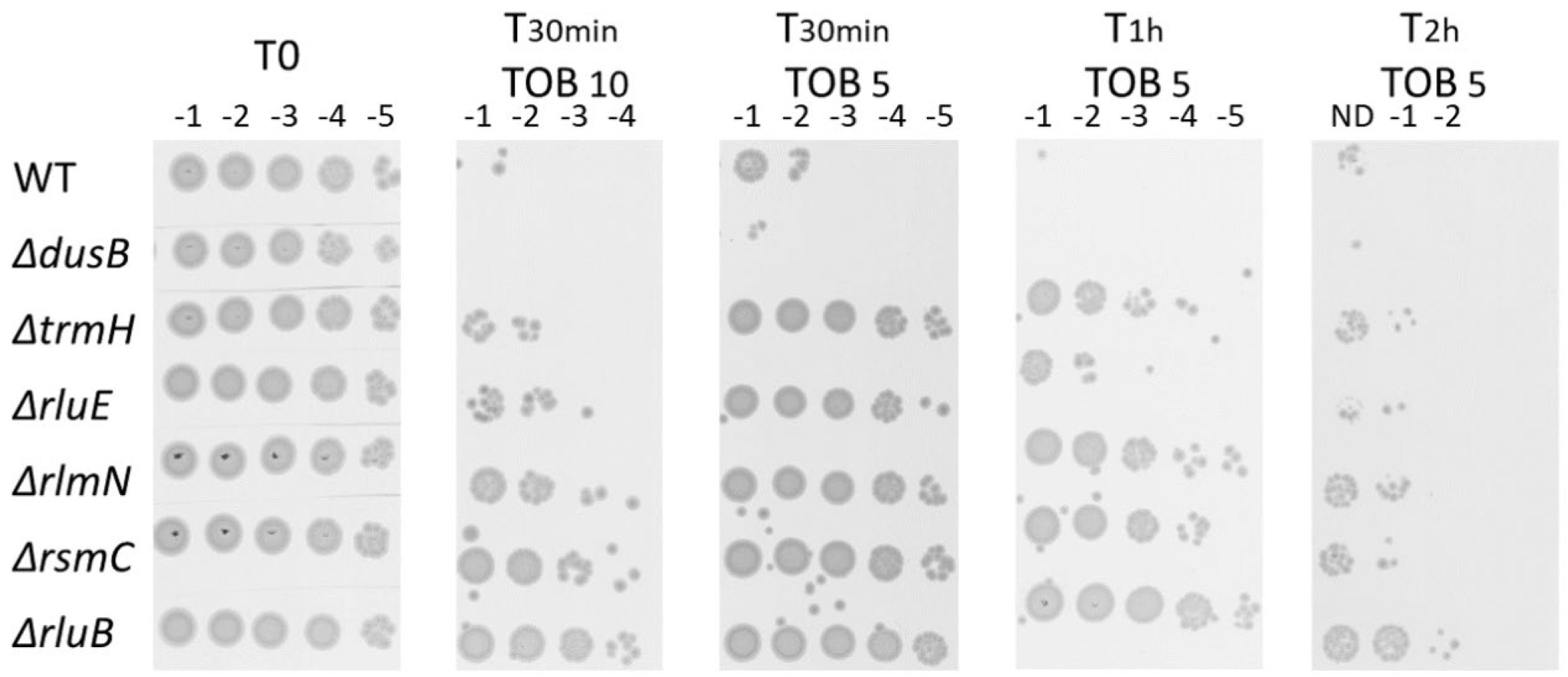
Survival to lethal antibiotic treatment. *V. cholerae* WT and deletion mutant cultures were grown without antibiotics up to early exponential phase, and serial dilutions were spotted on MH medium without antibiotics. Exponential phase cultures were then treated with antibiotics at lethal concentrations for 30min, 1h and 2 hours. At each time point, dilutions were spotted on MH. TOB: tobramycin 5 or 10 µg/ml.

**Figure S2.**
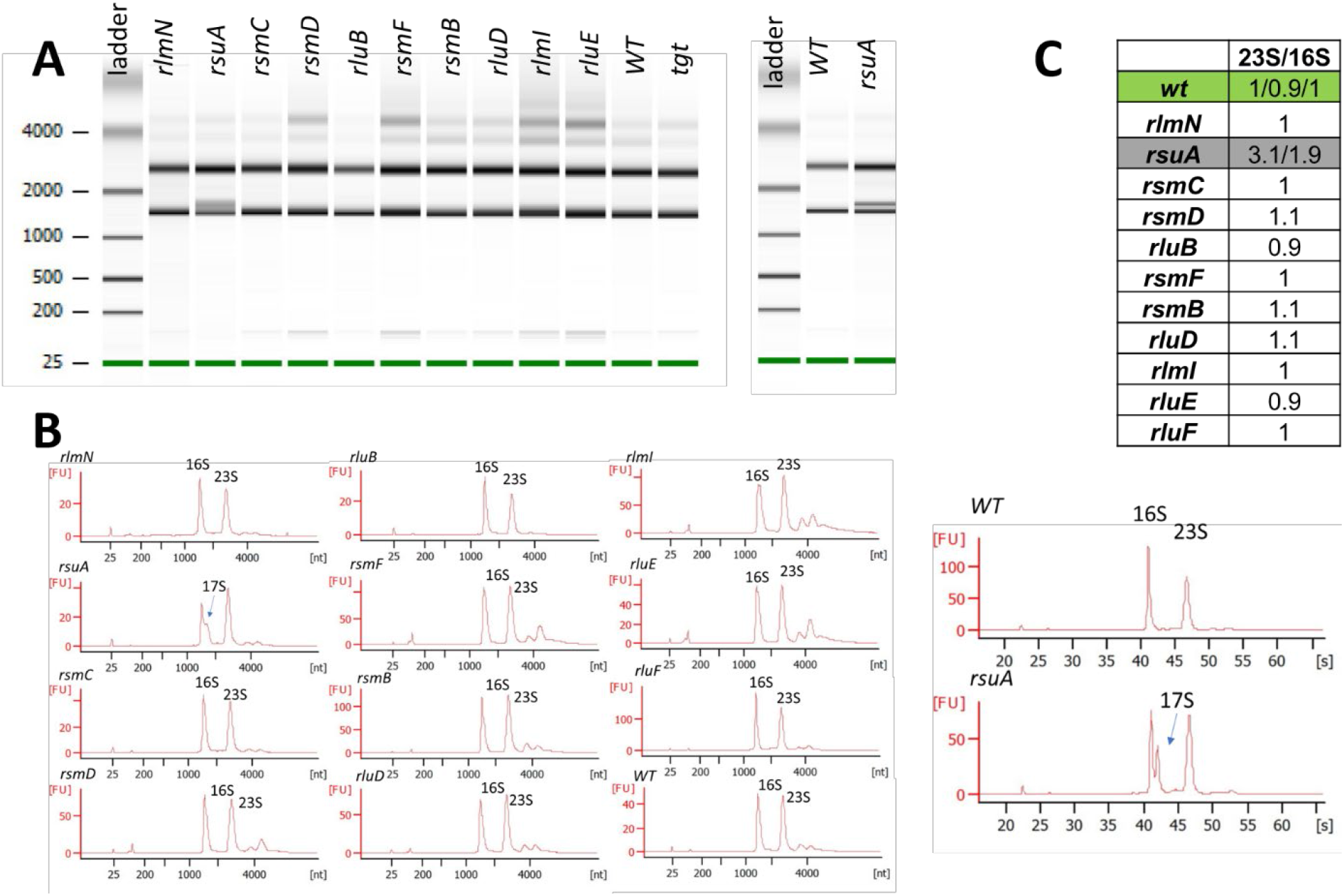
Effect of rRNA modification gene deletions on detected rRNA species during growth in the absence of antibiotics. A and B. Bioanalyzer results. C : 23S/16S ratio

**Figure S3.**
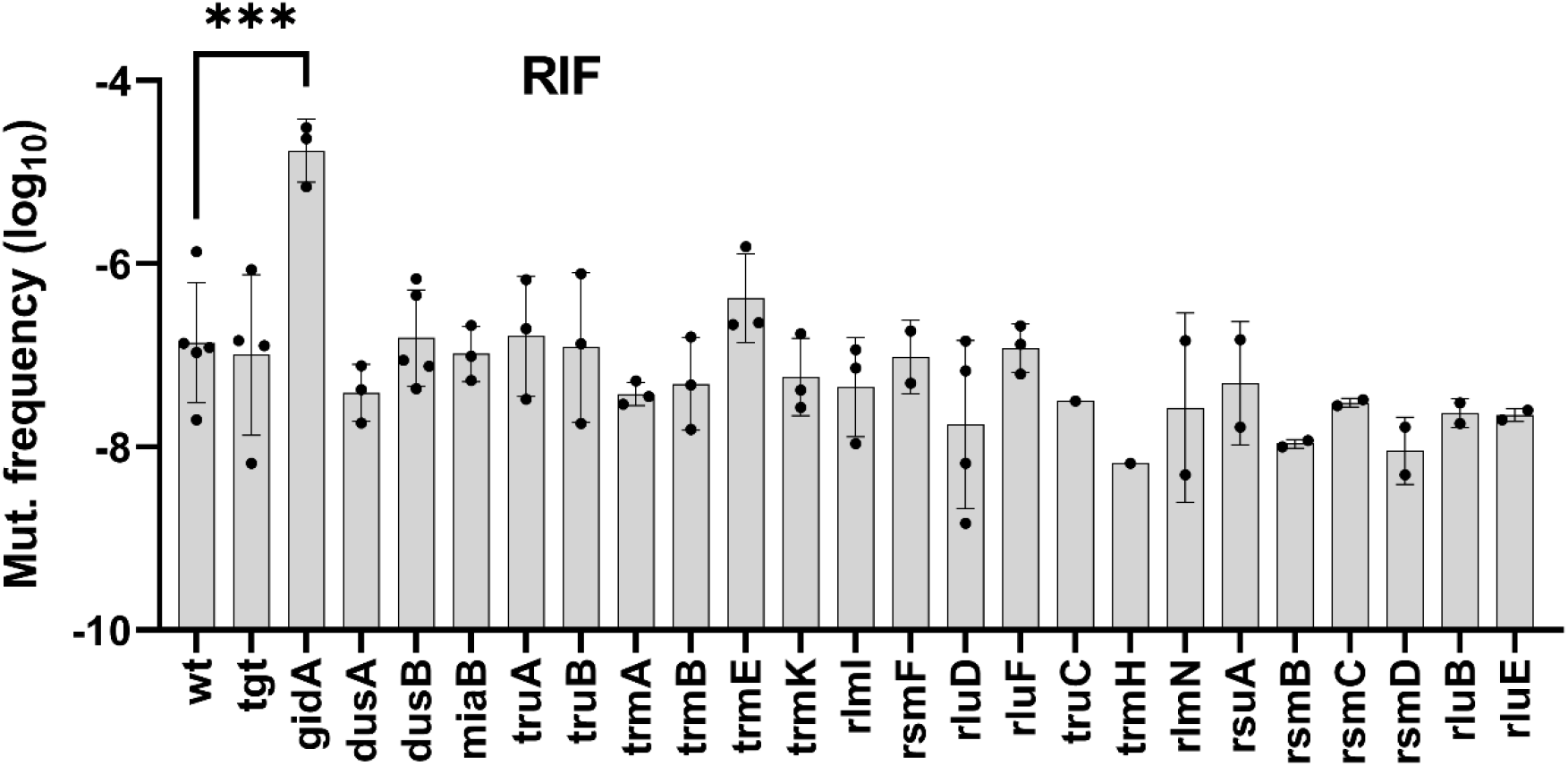
Frequency of appearance of spontaneous mutants in *V. cholerae* WT and RNA modification deletion mutants. Stationary phase cultures were plated in parallel on MH and MH plate supplemented with RIF: rifampicin 1 µg/ml. The mutation frequency was calculated as CFU MH + RIF/total CFU on MH. The Y-axis represents the log10 resistant mutant frequency. Data were first log transformed in order to achieve normal distribution, and statistical tests were performed on these log-transformed data. Means and geometric means for logarithmic values were calculated using GraphPad Prism. For multiple comparisons, we used one-way ANOVA on GraphPad Prism. **** means p<0.0001, *** means p<0.001. Number of replicates for each experiment: 3<n<8.

**Figure S4.**
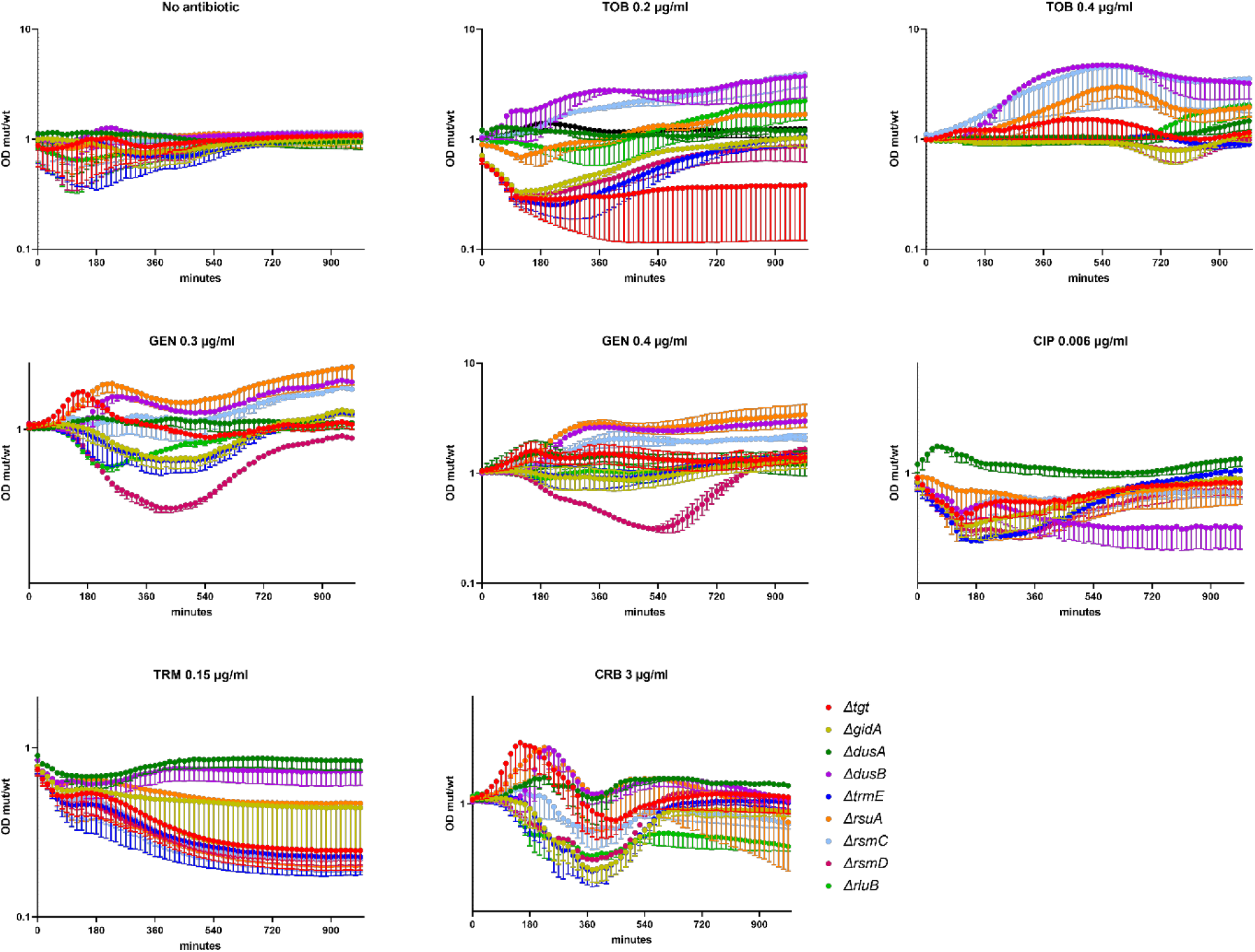
Growth of *E. coli* MG1655 WT and derivatives deleted for selected RNA modification genes. Overnight cultures were diluted 100x in fresh MH medium, on 96 well plates. Each well contained 200 µl. Plates were incubated with shaking on TECAN plate reader device at 37°C, OD 620 nm was measured every 15 minutes. Antibiotics were used at sub-MIC for MG1655 in MH liquid culture: TOB 0.2 and 0.4 µg/ml, CRB 3 µg/ml, TRM 0.15 µg/ml, GEN 0.3 and 0.4 µg/ml, CIP 0.006 µg/ml. The Y-axis represents the OD of the mutant grown in a given antibiotic divided by the OD of the WT strain in the same antibiotic, and thus reflects slower (below 1) or faster (above 1) growth. The Y-axis represents the OD of the mutant divided by the OD of the WT strain in the same growth conditions, and thus reflects slower (below 1) or faster (above 1) growth.

**Figure S5.**
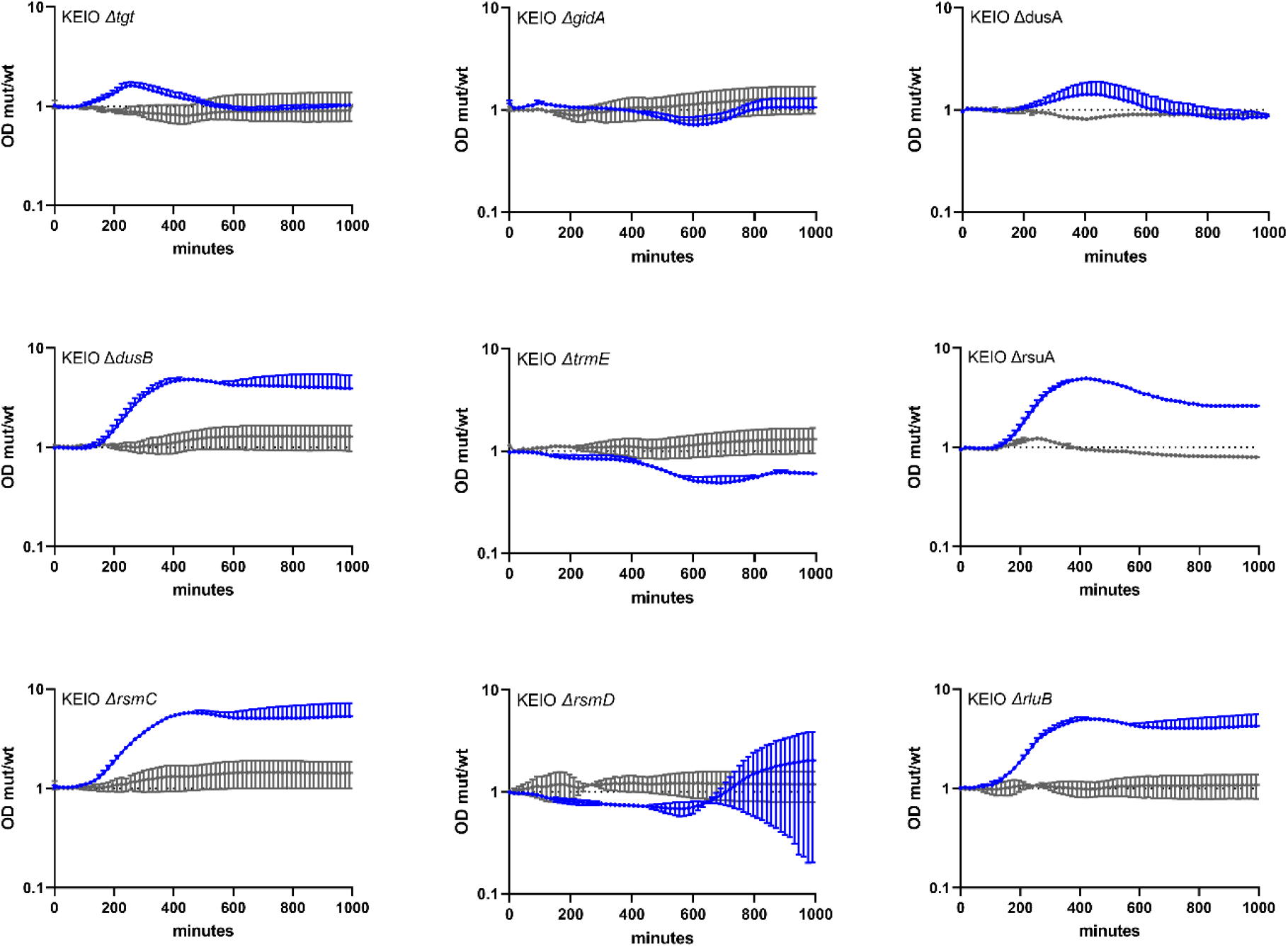
Growth of *E. coli* BW25113 WT and derivatives deleted for selected RNA modification genes in sub-MIC TOB. Overnight cultures were diluted 100x in fresh MH medium, on 96 well plates. Each well contained 200 µl. Plates were incubated with shaking on TECAN plate reader device at 37°C, OD 620 nm was measured every 15 minutes. Grey: no treatment. Blue: sub-MIC TOB, at 0.2 µg/ml (50% of the MIC for MG1655 in MH liquid culture). The Y-axis represents the OD of the mutant grown in a given antibiotic divided by the OD of the WT strain in the same growth conditions, and thus reflects slower (below 1) or faster (above 1) growth.

## Mat & Met

### Bacterial Strains and Plasmids

All *V. cholerae* strains used in this study are derivative of *V. cholerae* N16961 *hapR*+, and were constructed by allelic exchange. All *V. cholerae* mutants were constructed in the *ΔlacZ* strain (K329). All *E. coli* strains used in this work are derivatives of *E. coli* MG1655, and were constructed by transduction using *E. coli* Keio knockouts strains. Strains and plasmids are listed in **Tables S3 and S4** for more details.

### Media and Growth Conditions

Colonies on plates grew at 37°C, in Mueller-Hinton medium (MH) media. Liquid cultures grew at 37°C in appropriate media in aerobic conditions, with 180 rotations per minute

### Transposon insertion sequencing

**L**ibraries were prepared as previously described ^100,101^. to achieve a library size of 600.000 clones, and subjected to passaging in MH and MH+TOB 0.5 or MH+CIP 0,001 for 16 generations^33^. A saturated mariner mutant library was generated by conjugation of plasmid pSC1819 from *E.coli* to *V. cholerae* WT. Briefly, pSC189^100,101^ was delivered from *E. coli* strain 7257 (β2163 pSC189::spec, laboratory collection) into the *V. cholerae* WT strain. Conjugation was performed for 2 h on 0.45 µM filters. The filter was resuspended in 2 ml of MH broth. Petri dishes containing 100 µg/ml spectinomycin were then spread. The colonies were scraped and resuspended in 2 ml of MH. When sufficient single mutants were obtained (>600 000 for 6X coverage of non-essential regions), a portion of the library was used for gDNA extraction using Qiagen DNeasy® Blood & Tissue Kit as per manufacturer’s instructions. This was used for library validation through insert amplification by nested PCR using a degenerate primer (ARB6), which contains 20 defined nucleotides followed by a randomized sequence. This was combined with a primer anchored in the edge of the transposon sequence (MV288)^33,100^. After this, primer ARB3, which contains the first 20 nucleotides of ARB6 was used for nested amplification in combination with MV288. After validation, the libraries were passaged in MH media for 16 generations with or without 50%MIC of TOB or CIP, in triplicate. gDNA from time point 0 and both conditions after 16 generation passage in triplicate was extracted. Sequencing libraries were prepared using Agilent’s sureselect XT2 Kit with custom RNA baits designed to hybridize the edges of the Mariner transposon. The 100 ng protocol was followed as per manufacturer’s instructions. A total of 12 cycles were used for library amplification. Agilent’s 2100 bioanalzyer was used to verify the size of the pooled libraries and their concentration. HiSeq Paired-end Illumina sequencing technology was used producing 2×125bp long reads. Reads were then filtered through transposon mapping to ensure the presence of an informative transposon/genome junction using a previously described mapping algorithm^102^. Informative reads were extracted and mapped. Reads were counted when the junction was reported as mapped inside the CDS of a gene plus an additional 50 bp upstream and downstream. Expansion or decrease of fitness of mutants was calculated in fold changes with normalized insertion numbers. Normalization calculations were applied according to van Opijnen et al^103^. Expansion or decrease of fitness of mutants was calculated in fold changes with normalized insertion numbers. Baggerly’s test on proportions^104^ was used to determine statistical significance as well as a Bonferroni correction.

### Competitions experiments

Overnight cultures from single colonies of mutant lacZ+ and WT lacZ-strains were washed in PBS (Phosphate Buffer Saline) and mixed 1:1 (500μl + 500μl). At this point 100μl of the mix were serial diluted and plated on MH agar supplemented with X-gal (5-bromo-4-chloro-3-indolyl-β-D-galactopyranoside) at 40 μg/mL to assess T0 initial 1:1 ratio. At the same time, 10 μl from the mix were added to 2 mL of MH or MH supplemented with sub-MIC antibiotics (TCL: triclosan 0.01 µM, TOB: tobramycin 0.6 μg/ml; GEN: gentamicin 0.5 μg/ml; CIP: ciprofloxacin 0.002 µg/ml, CM: chloramphenicol 0.4 µg/ml, TRM: trimethoprim 0.4 µg/ml, CRB: carbenicillin 2.5 µg/ml) and incubated with agitation at 37°C for 20 hours. Cultures were then diluted and plated on MH agar plates supplemented with X-gal. Plates were incubated overnight at 37°C and the number of blue and white CFUs was assessed. Competitive index was calculated by dividing the number of blue CFUs (lacZ+ strain) by the number of white CFUs (lacZ-strain) and normalizing this ratio to the T0 initial ratio.

### MIC determination

Stationary phase cultures were diluted 20 times in PBS, and 300 µL were plated on MH plates and dried for 10 minutes. Etests (Biomérieux) were placed on the plates and incubated overnight at 37°C.

### Quantification and statistical analysis

First an F-test was performed in order to determine whether variances are equal or different between comparisons. For comparisons with equal variance, Student’s t-test was used. For comparisons with significantly different variances, we used Welch’s t-test. For multiple comparisons, we used one-way ANOVA. We used GraphPad Prism to determine the statistical differences (p value) between groups. **** means p<0.0001, *** means p<0.001, ** means p<0.01, * means p<0.05. For survival tests, data were first log transformed in order to achieve normal distribution, and statistical tests were performed on these log-transformed data. Number of replicates for each experiment was 3<n<6. Means and geometric means for logarithmic values were also calculated using GraphPad Prism. For persistence tests, data were first log transformed in order to achieve normal distribution, and statistical tests were performed on these log-transformed data.

### Survival/tolerance tests

were performed on early exponential phase cultures. In order to clear the culture from previously non-growing cells that could potentially be present from the stationary phase inoculum, we performed a two-step dilution protocol, before antibiotic treatment. Overnight *V. cholerae* cultures were first diluted 1000x in 4 ml fresh Mueller-Hinton (MH) medium, and incubated at 37°C with shaking. When the OD 620 nm reached ∼0.2, cultures were diluted 1000x a second time, in order to clear them from non-growing cells, in Erlenmeyers containing 25 ml fresh MH medium, and were allowed to grow at 37°C. When cultures reached an OD 620 nm between 0.25 and 0.3 (early exponential phase), appropriate dilutions were plated on MH plates to determine the total number of CFUs in time zero untreated cultures. Note that for *V. cholerae*, it was important to treat cultures at the precise OD 620 nm 0.25-0.3, as persistence levels seem to be particularly sensitive to growth phase in this species, where they decline in stationary phase. 5 ml of cultures were collected into 50 ml Falcon tubes and treated with lethal doses of desired antibiotics (10 times the MIC: tobramycin 10 µg/ml, carbenicillin 50 µg/ml, ciprofloxacin 0.025 µg/ml, trimethoprim 5 µg/ml) for 20 hours at 37°C with shaking in order to guarantee oxygenation. Appropriate dilutions were then plated on MH agar without antibiotics and proportion of growing CFUs were calculated by doing a ratio with total CFUs at time zero. Experiments were performed 3 to 8 times.

### RNA purification and analysis of rRNA species

For RNA extraction, overnight cultures were diluted 1:1000 in MH medium and grown with agitation at 37°C until an OD600 of 0.3 (exponential phase). 0.5 mL of these cultures were centrifuged and supernatant removed. Pellets were homogenized by resuspension with 1.5 mL of cold TRIzol Reagent. Next, 300 μL chloroform were added to the samples following mix by vortexing. Samples were then centrifuged at 4°C for 10 minutes. Upper (aqueous) phase was transferred to a new 2mL tube and mixed with 1 volume of 70% ethanol. From this point, the homogenate was loaded into a RNeasy Mini kit (Qiagen) column and RNA purification proceeded according to the manufacturer’s instructions. Samples were then subjected to DNase treatment using TURBO DNA-free Kit (Ambion) according to the manufacturer’s instructions. Total RNA samples were then analyzed on an Agilent 2100 Bioanalyzer (Agilent Technologies) using the Agilent RNA 6000 nano kit according to the instructions of the manufacturer.

### Rifampicin spontaneous mutation tests

Stationary phase cultures were plated in parallel on MH and MH plate supplemented with RIF: rifampicin 1 µg/ml. The mutation frequency was calculated as CFU MH + RIF/total CFU on MH.

### Growth of *E. coli* on microtiter plate reader

Overnight cultures were diluted 100x in fresh MH medium, on 96 well plates. Each well contained 200 µl. Plates were incubated with shaking on TECAN plate reader device at 37°C, OD 620 nm was measured every 15 minutes. Antibiotics were used at sub-MIC for MG1655 in MH liquid culture: TOB 0.2 and 0.4 µg/ml, CRB 3 µg/ml, TRM 0.15 µg/ml, GEN 0.3 and 0.4 µg/ml, CIP 0.006 µg/ml.

**Table S2.** Minimal inhibitory concentrations determined using etests.

| *V. cholerae* | TOB | CIP | AMP | TRM |
| --- | --- | --- | --- | --- |
| *WT* | 0.75 - 1.2 | 0.0020 - 0.0030 | 4 +/-1 | 0.4 +/- 0.1 |
| *tgt* | 0.75 - 1 | 0.0020 | 4 | 0.58 |
| *dusA* | 1 - 1 | 0.0020 - 0.0030 | 4 | 0.5 |
| *dusB* | 1 - 1 | 0.0020 - 0.0030 | 4 | 0.48 |
| *gidA* | 2 - 2 | 0.0039 | 4 | 0.37 |
| *trmA* | 0.9 - 1 | 0.0020 | 3.9 | 0.37 |
| *trmB* | 1 - 1.2 | 0.0020 | 4.3 | 0.48 |
| *miaB* | 0.9 - 1 | 0.0020 | 4 | 0.38 |
| *rsmF* | 1 | 0.0020 | 4.5 | 0.4 |
| *rlmI* | 1 | 0.0020 | 4.2 | 0.4 |
| *truA* | 0.8 | 0.0020 | 3.5 | 0.39 |
| *rluD* | 0.75 | 0.0036 | 4 | 0.5 |
| *trmE* | 0.75 - 1.2 | 0.0036 | 5 | 0.8 |
| *trmK* | 1 - 1.2 | 0.0020 | 4 | 0.38 |
| *rsmB* | 1 | 0.0020 | 4.1 | 0.48 |
| *truB* | 1 - 1.2 | 0.0020 | 4 | 0.47 |
| *rluF* | 1 - 1.2 | 0.0020 | 4 | 0.43 |
| *truC* | 0.75 | 0.0035 | 5 | 0.8 |
| *trmH* | 1 | 0.0025 | 4.3 | 0.5 |
| *rlmN* | 1 | 0.0025 | 5.5 | 0.38 |
| *rlmI* | 0.7 | 0.0028 | 4.5 | 0.44 |
| *rsuA* | 0.75 | 0.0030 | 3.5 | 0.4 |
| *rsmC* | 1.5 | 0.0028 | 4.8 | 0.38 |
| *rsmD* | 0.7 | 0.0028 | 3.5 | 0.38 |
| *rsmF* | 0.8 | 0.0032 | 3.8 | 0.38 |
| *rluB* | 1.5 | 0.0030 | 4 | 0.5 |
| *rluE* | 0.75 | 0.0028 | 5 | 0.5 |

**MIC determination using etests.** MH stationary phase cultures were diluted 20 times in PBS, and 300 µL were plated on MH plates and dried for 10 minutes. Etests (Biomérieux) were placed on the plates and incubated overnight at 37°C. **AMP:** ampicillin. This etest was used for carbenicillin evaluation. Green: increase MIC. Blue: decrease MIC.

**Table S3.** Strains and plasmids.

| Strain | Strain number | Construction |
| --- | --- | --- |
| *Vibrio cholerae* |  |  |
| N16961 hapR+ wt strain | F606 | Gift from Melanie Blokesch |
| N16961 hapR+ wt strain $\Delta lacZ$ | K329 | deletion of lacZ by plasmid integration and excision by sucrose counterselection as described |
| $\Delta tgt$ (VC0741) | J420 | PCR amplification of 500bp up and down regions of VC0741 using primers ZIP431/432 and ZIP433/434. PCR amplicification of aadA7 conferring spectinomycin resistance on pAM34 using ZB47/48. PCR assembly of the VC0741::spec fragment using ZIP431/434 and allelic exchange by natural transformation. |
| $\Delta tgt$ (VC0741) | M087 | allelic exchange by integration and excision of conjugative suicide plasmid pMP7 L910, replacing the gene with frt::kan::frt as described previously (Val et al PLoS Genetics 2012, Negro et al, mBio 2019) |
| $\Delta gidA$ (VC2775) | H244 | PCR amplification of 500bp up and down regions of VC2775 using primers ZIP316/317 and ZIP318/319. PCR amplicification of aadA7 conferring spectinomycin resistance on pAM34 using ZIP320/321. PCR assembly of the VC2775::spec fragment using ZIP316/319 and allelic exchange by natural transformation. |
| $\Delta dusA$ (VC0379) | L607 | allelic exchange by integration and excision of conjugative suicide plasmid pMP7 L024, replacing the gene with frt::kan::frt |
| $\Delta dusB$ (VC0291) | L606 | allelic exchange by integration and excision of conjugative suicide plasmid pMP7 L416, replacing the gene with frt::kan::frt |
| $\Delta miaB$ (VC0962) | K013 | (Negro et al, mBio 2019) |
| $\Delta truA$ (VC0999) | N095 | allelic exchange by integration and excision of conjugative suicide plasmid pMP7 970, replacing the gene with frt::kan::frt |
| *ΔtruB (VC0645)* | M562 | allelic exchange by integration and excision of conjugative suicide plasmid pMP7 M347, replacing the gene with frt::kan::frt |
| *ΔtruC (VC0888)* | P638 | allelic exchange by integration and excision of conjugative suicide plasmid pMP7 O651, replacing the gene with frt::kan::frt |
| *ΔtrmA (VC0154)* | M564 | allelic exchange by integration and excision of conjugative suicide plasmid pMP7 M423, replacing the gene with frt::kan::frt |
| *ΔtrmB (VC0453)* | M096 | allelic exchange by integration and excision of conjugative suicide plasmid pMP7 L974, replacing the gene with frt::kan::frt |
| *ΔtrmE (VC0003)* | H218 | PCR amplification of 500bp up and down regions of VC0003 using primers 1640/1641 and 1642/1643. PCR amplicification of aadA7 conferring spectinomycin resistance on pAM34 using 1644/1645. PCR assembly of the VC0003::spec fragment using 1640/1643 and allelic exchange by natural transformation. |
| *ΔtrmH (VC0803)* | Q062 | allelic exchange by integration and excision of conjugative suicide plasmid pMP7 P493, replacing the gene with frt::kan::frt |
| *ΔtrmK (VCA0634)* | K650 | allelic exchange by integration and excision of conjugative suicide plasmid pMP7 K440, replacing the gene with frt::kan::frt |
| *ΔrlmN (VC0757)* | M094 | allelic exchange by integration and excision of conjugative suicide plasmid pMP7 L912, replacing the gene with frt::kan::frt |
| *ΔrlmI (VC1354)* | N031 | allelic exchange by integration and excision of conjugative suicide plasmid pMP7 M969, replacing the gene with frt::kan::frt |
| *ΔrsuA (VC1635)* | H497 | Negro et al, mBio 2019 |
| *ΔrsmB (VC0044)* | N033 | allelic exchange by integration and excision of conjugative suicide plasmid pMP7 M771, replacing the gene with frt::kan::frt |
| *ΔrsmC (VC0623)* | L601 | allelic exchange by integration and excision of conjugative suicide plasmid pMP7 L577, replacing the gene with frt::kan::frt |
| *ΔrsmD (VC0146)* | M088 | allelic exchange by integration and excision of conjugative suicide plasmid pMP7 L565, replacing the gene with frt::kan::frt |
| *ΔrsmF (VC2223)* | N045 | allelic exchange by integration and excision of conjugative suicide plasmid pMP7 M769, replacing the gene with frt::kan::frt |
| *ΔrluB (VC1179)* | L559 | allelic exchange by integration and excision of conjugative suicide plasmid pMP7 L020, replacing the gene with frt::kan::frt |
| *ΔrluD (VC0709)* | N097 | allelic exchange by integration and excision of conjugative suicide plasmid pMP7 N035, replacing the gene with frt::kan::frt |
| *ΔrluE (VC1140)* | Q061 | allelic exchange by integration and excision of conjugative suicide plasmid pMP7 P346, replacing the gene with frt::kan::frt |
| ***Escherichia coli*** |  |  |
| MG1655 wt strain |  | laboratory collection |
| *Δtgt* | J233 | P1 transduction from KEIO strain JW0396-3 |
| *ΔgidA* | J193 | P1 transduction from KEIO strain JW3719-1 |
| *ΔdusA* | J196 | P1 transduction from KEIO strain JW5950-5 |
| *ΔdusB* | J243 | P1 transduction from KEIO strain JW3228-1 |
| *ΔtrmE* | J194 | P1 transduction from KEIO strain JW3684-1 |
| *ΔrsuA* | H243 | P1 transduction from KEIO strain JW2171-1 |
| *ΔrsmC* | J192 | P1 transduction from KEIO strain JW4333-1 |
| *ΔrsmD* | J241 | P1 transduction from KEIO strain JW3430-4 |
| *ΔrluB* | J235 | P1 transduction from KEIO strain JW1261-3 |
| **Plasmids pMP7-Δgene::kan** |  | gibson assembly using primers MV450/451 for the amplification of pMP7 vector, primers indicated below for up and down regions of the gene, and primers MV268/269 on pKD4 plasmid for the resistance gene (frt::kan::frt). |
| *Δtgt (VC0741)* | L910 | VC0741tgt5/7 for up region and VC0741tgt6bis/8 bis for down region |
| *ΔdusA (VC0379)* | L024 | VC0379dusA5/7 for up region and VC0379dusA6/8 for down region |
| *ΔdusB (VC0291)* | L416 | VC0291dusB5/7 for up region and VC0291dusB6bis/8 bis for down region |
| *ΔtruA (VC0999)* | M970 | VC0999truA5/7 for up region and VC0999truA6bis/8bis for down region |
| *ΔtruB (VC0645)* | M347 | VC0645truB5bis/7bis for up region and VC0645truB6/8 for down region |
| *ΔtruC (VC0888)* | O651 | VC0888truC5/7 for up region and VC0888truC6/8 for down region |
| *ΔtrmA (VC0154)* | M423 | VC0154trmA5/7 for up region and VC0154trmA6bis/8 bis for down region |
| *ΔtrmB (VC0453)* | L974 | VC0453trmB5/7 for up region and VC0453trmB6/8 for down region |
| *ΔtrmH (VC0803)* | P493 | VC0803trmH5/7 for up region and VC0803trmH6bis/8 for down region |
| *ΔtrmK (VCA0634)* | K440 | VCA06345/7 for up region and VCA06346/8 for down region |
| *ΔrlmN (VC0757)* | L912 | VC0757rlmN5/7 for up region and VC0757rlmN6/8 for down region |
| *ΔrlmI (VC1354)* | M969 | VC1354rlmI5bis/7bis for up region and VC1354rlmI6/8 for down region |
| *ΔrsmB (VC0044)* | M771 | VC0044rsmB5/7 for up region and VC0044rsmB6/8 for down region |
| *ΔrsmC (VC0623)* | L577 | VC0623rsmC5/7 for up region and VC0623rsmC6/8 for down region |
| *ΔrsmD (VC0146)* | L565 | VC0146rsmD5/7 for up region and VC0146rsmD6/8 for down region |
| *ΔrsmF (VC2223)* | M769 | VC1502rsmF5/7 for up region and VC1502rsmF6/8 for down region |
| *ΔrluB (VC1179)* | L020 | VC1179rluB5/7 for up region and VC1179rluB6/8 for down region |
| *ΔrluD (VC0709)* | N035 | VC0709rluD5ter/7ter for up region and VC0709rluD6bis/8ter for down region |
| *ΔrluE (VC1140)* | P346 | VC1140rluE5/7 for up region and VC1140rluE6/8 for down region |

**Table S4.**
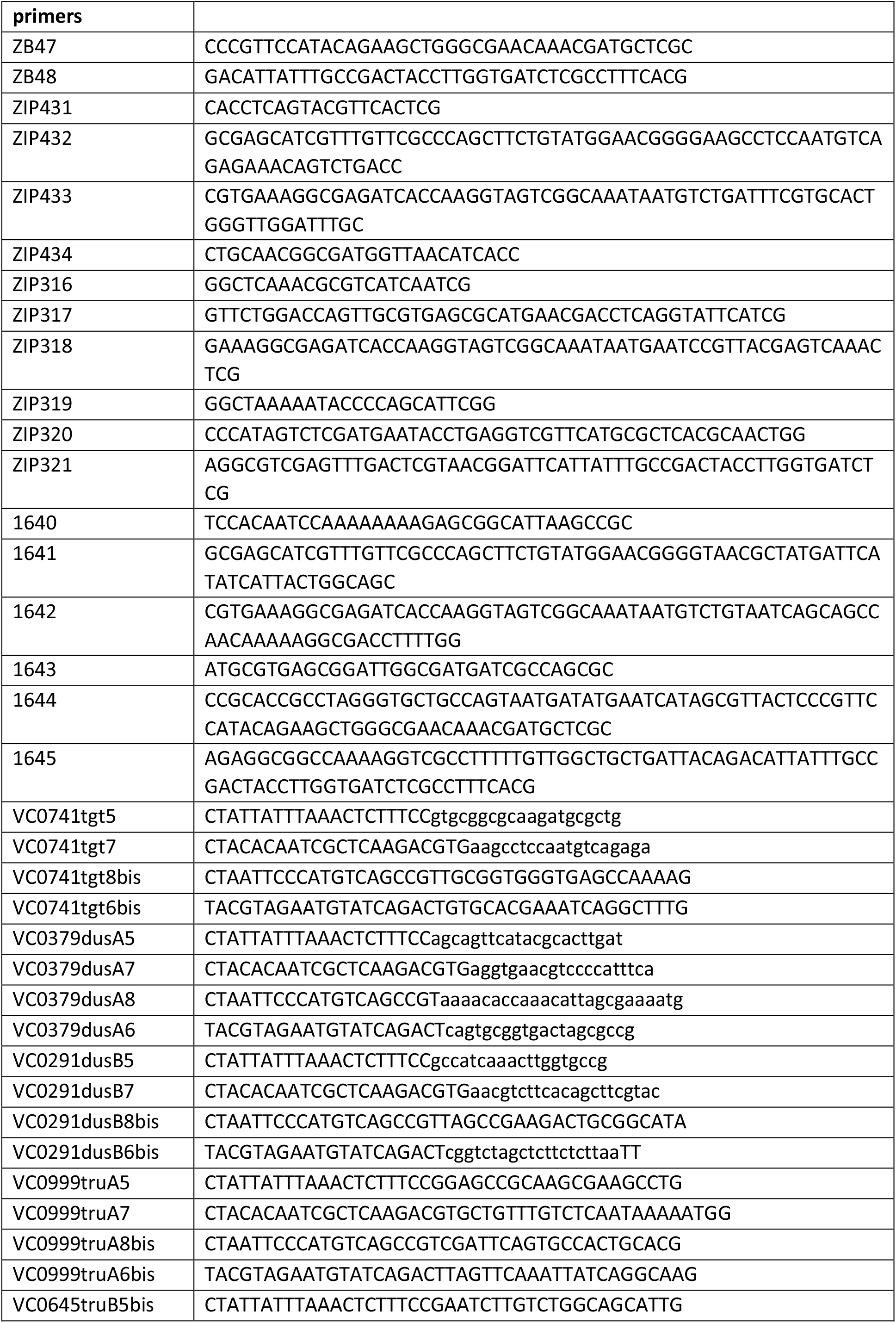
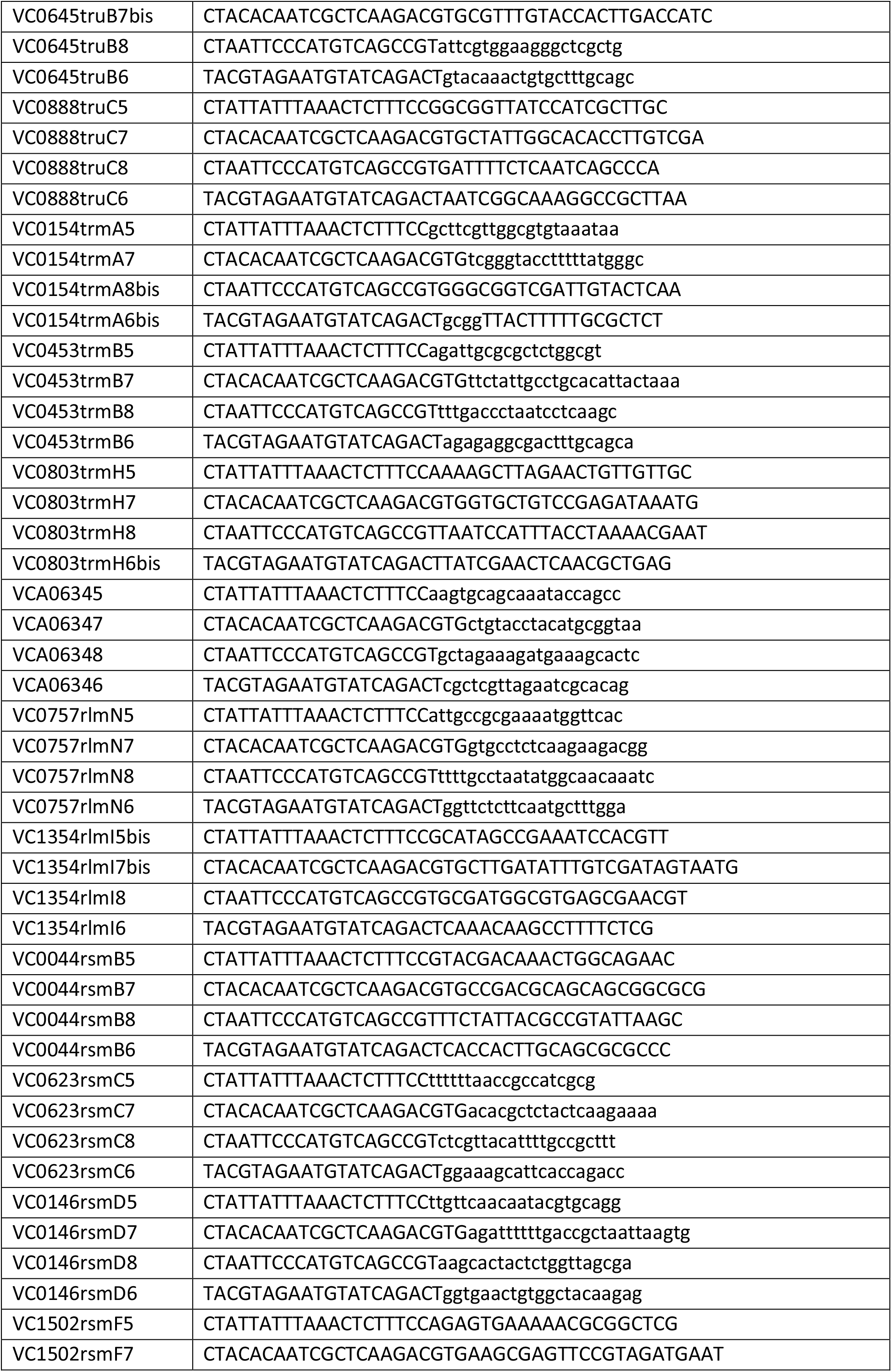
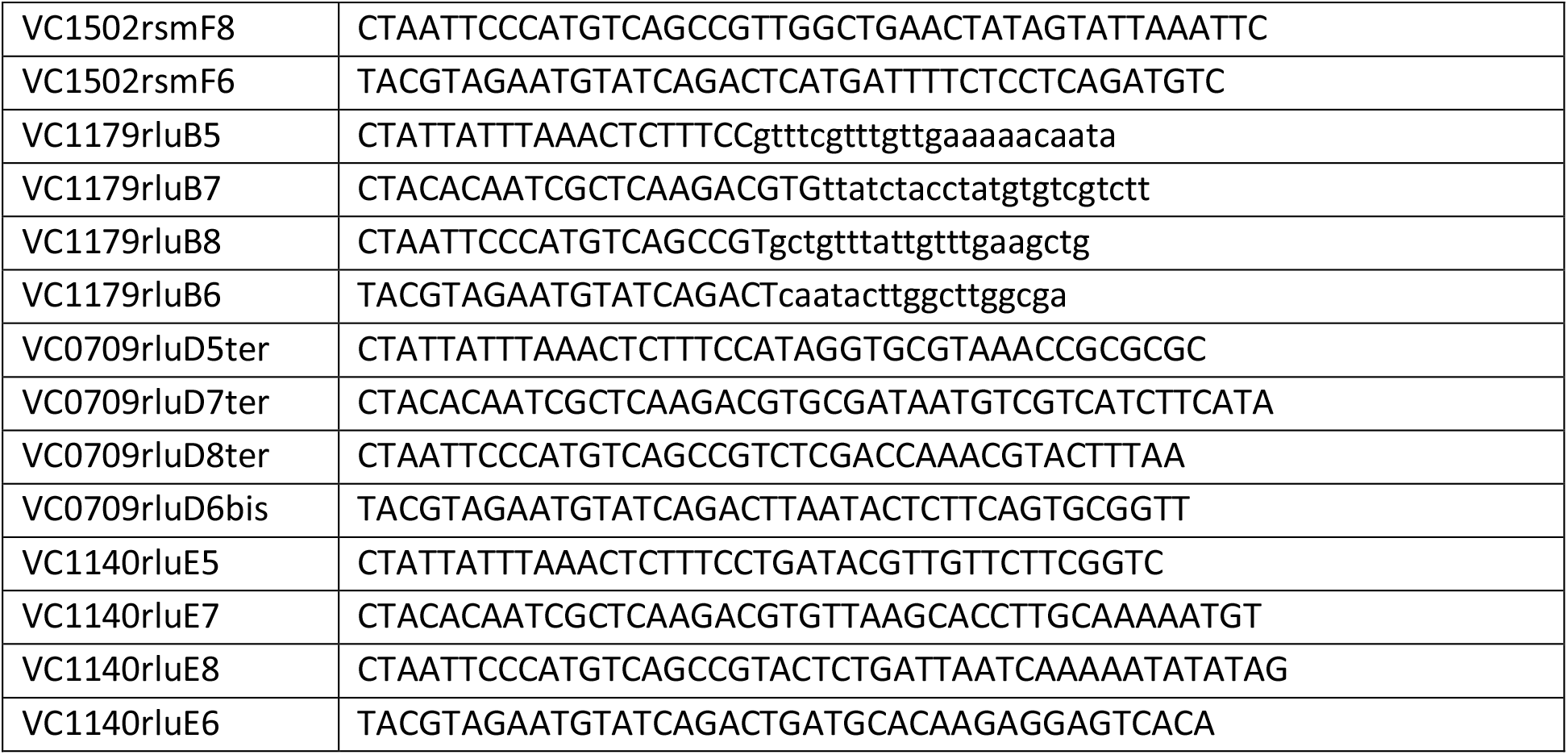
Primers.

